# dictyExpress: an integrated browser for bulk and single-cell *Dictyostelium* transcriptomics

**DOI:** 10.64898/2026.09.04.748563

**Authors:** Lena Trnovec, Miha Štajdohar, Gad Shaulsky, Blaž Zupan

## Abstract

**Summary:** dictyExpress 3.0 is a React/TypeScript reimplementation of the *Dictyostelium discoideum* transcriptomics web server, first introduced in 2009, that unifies bulk and single-cell exploration. A long-standing gene-expression resource for the *Dictyostelium* community, dictyExpress is also accessible through dictyBase, the central organism database for *Dictyostelium*. The new release preserves the curated bulk RNA-seq dashboards of earlier versions while adding an interactive module for a developmental scRNA-seq time course of wild-type AX4 and two cAMP-signalling mutants. Following visual analytics principles, interactive visualizations are coordinated through brushing and linking, with bulk time courses, differential expression, gene-ontology enrichment and the UMAP workspace sharing a common gene selection, enabling bookmarkable and exportable cross-modal analysis in one interface for *Dictyostelium* transcriptomics.

**Availability and implementation:** dictyExpress is available at https://app.dictyexpress.org/.Source code: https://github.com/biolab/dictyexpress-js (Apache License 2.0), archived at https://doi.org/10.5281/zenodo.22042272.

## Introduction

The social amoeba *Dictyostelium discoideum* is a model for multicellular development, cell motility and cAMP-mediated signalling (Eichinger et al., 2005; Williams, 2010; Loomis, 2014). Its 24-hour developmental programme, from solitary amoebae to a multicellular fruiting body, has been profiled extensively by bulk RNA-seq, producing developmental time courses, mutant comparisons and perturbation datasets that together anchor current understanding of gene regulation during this transition (Rosengarten et al., 2015; Katoh-Kurasawa et al., 2021). Single-cell RNA-seq (scRNA-seq) has since begun to resolve the cell-type heterogeneity that is averaged out in those population-level measurements, and a recent developmental time course of wild-type and cAMP-signalling mutants has placed such data at the centre of the *Dictyostelium* toolkit (Katoh-Kurasawa et al., 2026b). Earlier dictyExpress releases, however, were built around curated bulk datasets alone (Rot et al., 2009; Stajdohar et al., 2017), so this scRNA-seq resource had no integrated companion in the community’s primary exploration tool.

dictyExpress has served as one of the central *Dictyostelium* community tools since its first release. Version 1.0 introduced exploratory access to microarray gene-expression profiles (Rot et al., 2009), and version 2.0 extended the platform to RNA-seq data to support differential-expression analysis, hierarchical clustering, reporting of GO enrichment, and search for similarly expressed genes at client side through Resolwe-based dataflow back end engine (Stajdohar et al., 2017). Version 3.0 builds on these curated bulk data workflows, extends them to manage scRNA-seq data, and pairs them with a new web client whose client-side rendering and caching keep single-cell datasets of tens of thousands of cells responsive in the browser. The result is a unified reactive environment for bulk and single-cell exploration of *Dictyostelium* transcriptomes (Figure 1a), in which changes in settings or data selections in one visual panel propagate to the others through coordinated views (Roberts, 2007), thus supporting exploratory analysis and visual analytics.

**Figure 1.**
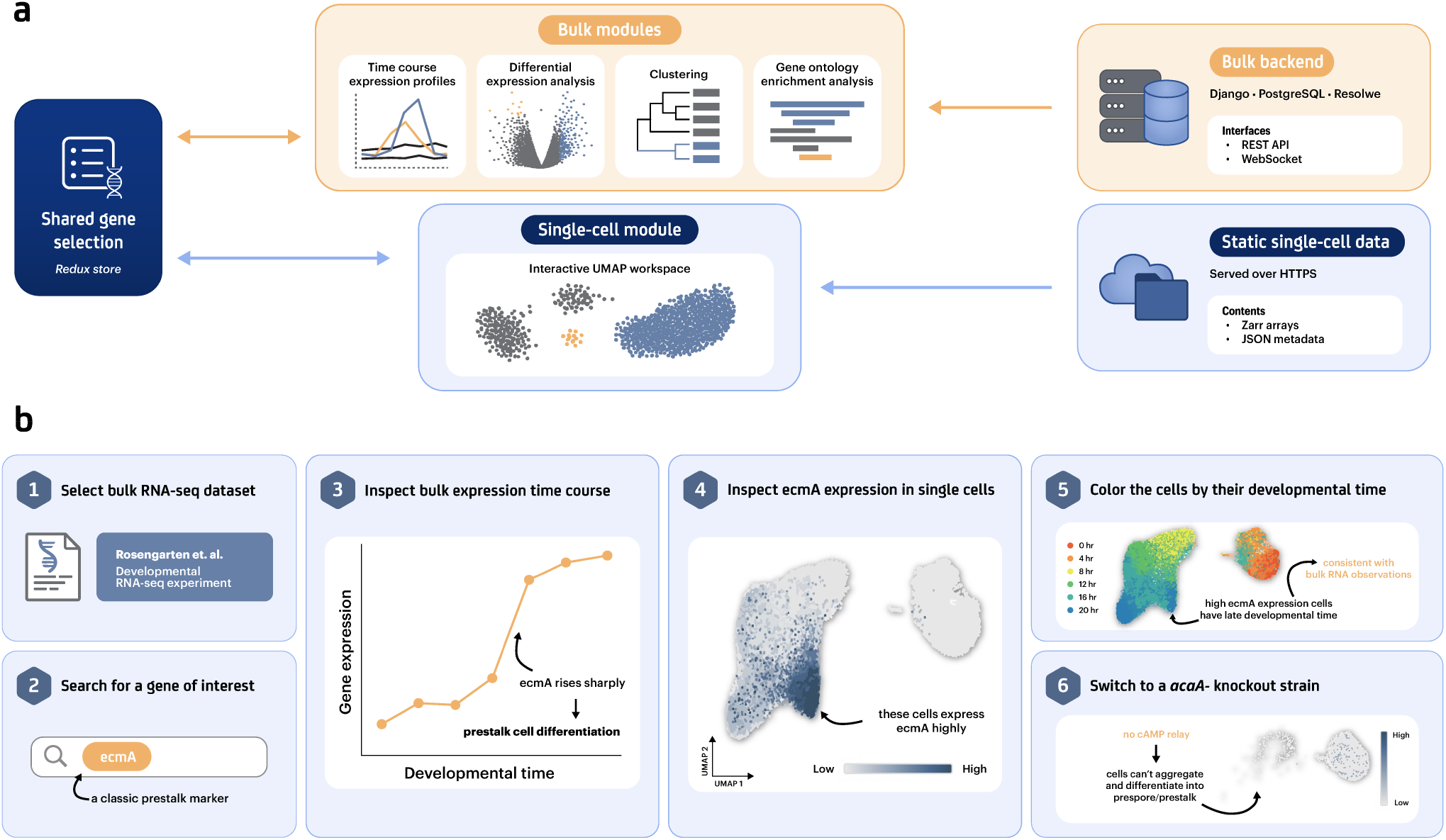
An overview of dictyExpress 3.0. **(a)** System architecture. A shared gene selection, held in a client-side Redux store, couples the bulk-analysis modules (expression time courses, differential expression, hierarchical clustering and gene-ontology enrichment; served by the Django / PostgreSQL / Resolwe back end) to the interactive single-cell UMAP workspace (served as static per-gene Zarr chunks with accompanying UMAP and annotation metadata). Selecting or refining a gene set in any module updates the store and propagates to all other modules. **(b)** Representative bulk-to-single-cell workflow. (1) The user loads a curated developmental bulk RNA-seq dataset (Rosengarten et al., 2015) and (2) searches for a marker gene such as *ecmA*. (3) Its developmental trajectory is inspected in the bulk time-course module, and (4) the same gene is followed directly into the single-cell UMAP, where each cell is coloured by its relative expression level. (5) Recolouring by developmental time (0–20 h) places the expression signal in its temporal context. (6) Switching to the *acaA*^−^ mutant, which lacks the ability to synthesize cAMP, preserves the AX4 reference coordinate frame, so wild-type and mutant patterns can be compared without re-running an analysis.

## Features

### Dashboard and bulk transcriptomics

dictyExpress 3.0 reimplements the core bulk RNA-seq workflow of v2.0 (Stajdohar et al., 2017) in a new React/TypeScript client while preserving its analytical behaviour. Users begin in an experiment browser that lists curated developmental time courses and perturbation studies, each linked to its source publication and annotated with strain, treatment and growth conditions. Selecting an experiment populates a drag-and-drop dashboard of six coordinated modules (Figure 1a): (1) *Time Series and Gene Selection*, with autocomplete gene search; (2) *Expression Time Courses*, plotting normalised-count trajectories and supporting similar-gene search and cross-dataset overlays; (3) *Differential Expression*, with volcano plots and an interactive selection window for filtering selected genes; (4) *Hierarchical Clustering* of genes based on expression profile similarity, supporting interactive brushing for gene selection, (5) *Gene Ontology Enrichment* for displaying of enriched GO terms of currently selected genes; and (6) *Single-Cell Expression*, the new interactive UMAP workspace and single-cell module described below.

A shared gene list drives and connects all six modules, allowing users to move between complementary views without re-entering the same query. Changes of gene list in one module are propagated to all other modules, refreshing their display. Data visualization modules become informative only after gene list has been selected. Gene lists can be saved and recalled from a history dialog, and each gene entry deep-links to dictyBase (Fey et al., 2019) for community annotation. Analysis sessions can be bookmarked as shareable URLs and exported as ZIP archives of publication-ready figures and tables; both operate on the whole session, covering the chosen datasets, genes, highlights, module settings and single-cell views, rather than on individual modules. The first five modules thus preserve the established bulk workflow from previous versions of dictyExpress, while the sixth extends it into single-cell analysis.

### Single-cell module

The central addition in v3.0 is an interactive single-cell UMAP workspace that hosts a recent developmental scRNA-seq time course (Katoh-Kurasawa et al., 2026b). The dataset spans three *D. discoideum* strains, namely wild-type AX4 and two cAMP-signalling mutants (*acaA*^−^ and *acaA*^−^*pkaCOE*), collected at six time points across 0–20 h and profiled with 10x Chromium scRNA-seq (Zheng et al., 2017). The AX4 dataset alone comprises 48,983 cells and 12,883 genes. Cells were embedded with the Universal Cell Embedding (UCE) foundation model (Rosen et al., 2023) and mapped to two dimensions with UMAP (McInnes et al., 2018; Becht et al., 2019). Mutant cells are mapped into the AX4 reference embedding so that all the strains share a single coordinate frame.

The UMAP visualization supports three colouring modes: (i) gene expression, with user-selectable aggregation (average, sum, minimum, or maximum when 2 or more genes are selected) and linear or log_1*p*_ transform; (ii) developmental time (0–20 h); and (iii) cell type. In the time and cell-type modes an optional expression-alpha overlay modulates point opacity by aggregated expression, so users can see how strongly a gene set is expressed within each time window or cell-type cluster. When multiple genes are selected (from other modules), UMAP visualization applies aggregation function per cell across the gene set, making pathway-level signatures directly visible on the embedding. The UMAP is rendered on an HTML5 Canvas, supports pan, zoom and double-click reset, and holds the AX4 viewport bounds fixed when users switch strains, so that point positions remain directly comparable across strains.

#### Bulk–single-cell integration

All six modules share a single gene selection through a centralised Redux store, and this shared state is the central design principle of dictyExpress 3.0. Choosing a gene in any module simultaneously plots its bulk normalised-count trajectory, colours every cell on the UMAP by that gene’s expression in each strain, and makes the same gene set available to differential-expression and GO-enrichment queries without reconfiguration. When the user highlights a subset of genes, downstream enrichment operates on that subset rather than on the full selection, so a broad candidate list can be narrowed to a more coherent transcriptional programme in a few clicks. The “find similar genes” function in the bulk time-course module feeds its co-expressed hits back into the same shared state, providing a direct path from bulk co-expression to single-cell spatial patterns.

#### Client-side data architecture

To keep the UMAP responsive, single-cell expression matrices are stored as per-strain Zarr arrays (Miles et al., 2020) chunked by gene (1 *× n*_cells_) and served as static files. Because each chunk corresponds to a single gene, querying expression for one gene transfers only that gene’s vector rather than the full matrix. The browser reads these arrays via the zarr JavaScript library and caches each gene slice in IndexedDB, and a shared HTTPStore instance amortises TCP/TLS connection setup across successive queries. The net effect is that UMAP colouring remains interactive without any server-side computation for single-cell rendering. Adding new sc-RNA datasets, as they become available, requires only deploying a strain-specific directory under `/single-cell-data/` containing the Zarr expression array together with JSON files for gene identifiers, cell annotations, UMAP coordinates and dataset metadata; available strains are read from a top-level `strains.json`. The current release serves one curated developmental scRNA-seq time course spanning the three strains above, but the layout is designed to accommodate additional strains or conditions without any change to the application code.

### Representative analysis workflow

dictyExpress 3.0 is designed for exploratory analysis and cross-checking between bulk and single-cell views, not for upstream scRNA-seq processing (Figure 1b). A typical session begins by selecting a curated bulk dataset, such as the Rosengarten developmental time course (Rosengarten et al., 2015). The user searches for a marker, inspects its expression trajectory, and follows the same gene directly into the single-cell UMAP. From there, the cells can be recoloured by developmental time or by cell type to place the signal in context, and the view can be switched to a mutant strain within the same coordinate frame.

User’s session can also branch into differential expression. Genes brushed from the volcano plot enter the shared gene set, where they can be clustered by temporal behaviour and summarised by GO enrichment. Highlighting a dendrogram branch or a subset of genes in the shared selection narrows enrichment to that focused subset rather than the full set.

Because bookmarking and export capture that shared state in full, the whole workflow can be saved, reopened and shared. A 19-step in-app tutorial walks first-time users through this progression end to end, from marker search through bookmarking and export, and serves as a short guided case study of the reproducible bulk-to-single-cell workflow described above.

### Reproducibility and usability

Sharing a whole session in this way removes the usual reconstruction of an analysis from a written protocol, and the surrounding usability is tuned to match the same exploratory mode: the dashboard layout can be rearranged and persisted locally, letting each user emphasise the modules most relevant to a given exploration while keeping the underlying gene selection shared, and the in-app tutorial gives occasional users a concrete entry point into the workflow. Together with the public web deployment and browser-side handling of the single-cell matrices, these choices make dictyExpress 3.0 suitable both for rapid hypothesis generation and for lightweight sharing of analysis states between collaborators.

### Implementation

dictyExpress 3.0 is a multi-tier web application. The React/TypeScript front end provides the responsive drag-and-drop dashboard, persistent layouts and canvas-based single-cell rendering, with bulk-module visualisations built on Vega (Satyanarayan et al., 2017) and AG Grid. The back end (Django, PostgreSQL and the Resolwe dataflow engine (Stajdohar et al., 2017)) is unchanged from v2.0, and the front end communicates with it through reverse-proxied REST and WebSocket endpoints.

Two operations that users invoke repeatedly during interactive exploration, namely similar-gene search in the Expression Time Courses module and Gene Ontology enrichment, were originally implemented as Resolwe processes, so each request was scheduled in the worker queue and returned only after process start-up and an intermediate storage fetch. In v3.0 both can instead be answered by lightweight, stateless serverless endpoints (`/find-similar and /go`), deployed as AWS Lambda functions and exposed as same-origin routes on the application’s content-delivery distribution. The find-similar endpoint receives the selected time series, samples, query gene and distance metric and ranks matches directly against a precomputed, replicate-averaged time-course matrix, collapsing the original multi-step dataflow (job scheduling, process execution and a separate storage download) into a single same-origin request. The enrichment endpoint takes the selected genes together with their annotation context (source, species and ontology). In practice this removes the worker-queue and storage-transfer overhead that dominated the Resolwe path: for the developmental time course (Rosengarten et al., 2015), ranking the full set of ~12,000 genes took about 20 ms of server-side computation, against roughly 1.8 s previously spent merely downloading the result of the scheduled process.

The `/find-similar` and `/go` endpoints are configuration-gated: when they are not set, the two modules transparently fall back to their Resolwe processes, so the optimisation is purely additive and the client remains fully functional against a plain Resolwe back end. Single-cell data bypass the back end entirely and are served as static Zarr arrays over HTTPS, so the production build is self-contained and can be deployed from any HTTPS-capable web server: in production, `/api` and `/ws` requests are proxied to Resolwe, `/find-similar` and /go resolve to their Lambda functions, and the single-cell path serves static files directly.

## Conclusion

dictyExpress 3.0 brings single-cell transcriptomics into the same interface that the *Dictyostelium* community has used for bulk RNA-seq analysis since version 1.0 (Rot et al., 2009; Stajdohar et al., 2017). By connecting an interactive UMAP workspace to all previously established modules, the app lets users move between population-level and single-cell views of the same genes without switching tools or re-uploading data. The bookmarkable sessions, exportable figures and 19-step in-app tutorial together lower the barrier to adoption. The current release centres on a curated developmental scRNA-seq time course of AX4 and cAMP-signalling mutants, while the static Zarr layout supports adding data on new strains or conditions without changing the application code. The front end is open-source (Apache License 2.0) and runs in all modern browsers with no server-side session state, so it should be straightforward to maintain and extend as further *Dictyostelium* scRNA-seq datasets become available.

## Acknowledgements

We thank the Shaulsky laboratory at Baylor College of Medicine for generating the single-cell datasets and the Genialis team for maintaining the Resolwe back-end infrastructure.

## Data availability

dictyExpress 3.0 can be accessed at https://app.dictyexpress.org/. Its source code is available at https://github.com/biolab/dictyexpress-js under the Apache License 2.0. The application provides access to curated bulk transcriptomic datasets and to the developmental single-cell dataset reported by Katoh-Kurasawa et al. (2026b), which is available from the NCBI Gene Expression Omnibus under accession number GSE305468 (Katoh-Kurasawa et al., 2026a).

