## Supplementary Information for "dictyExpress: an integrated browser for bulk and single-cell *Dictyostelium* transcriptomics"

#### Use case from the built-in tutorial

Figure S1 shows steps 1–8 of the built-in dictyExpress tutorial. Using *ecmA*, a classic prestalk marker, it guides users from the "Filter Development vs. cAMP Pulsing" bulk dataset to the AX4 single-cell map and then colours cells by collection time. It next switches to the *acaA*- mutant, in which adenylyl cyclase A is deleted and cAMP relay is disrupted, to compare strains in the shared AX4 reference embedding. Later steps extend the shared selection to differential expression, gene selection, clustering, Gene Ontology enrichment, and export.

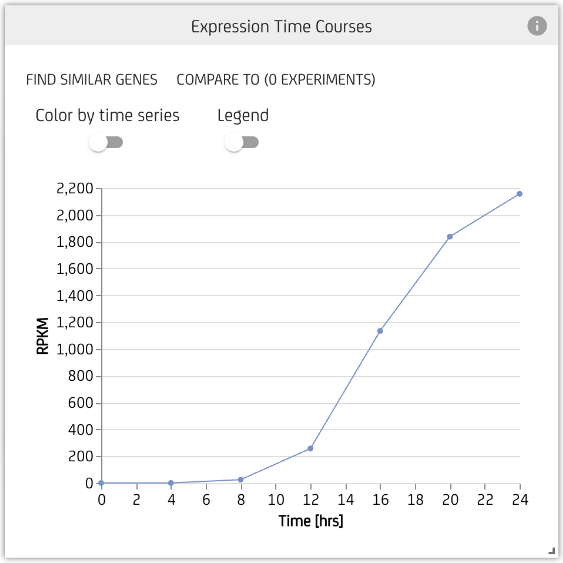


**A** Bulk *ecmA* time course

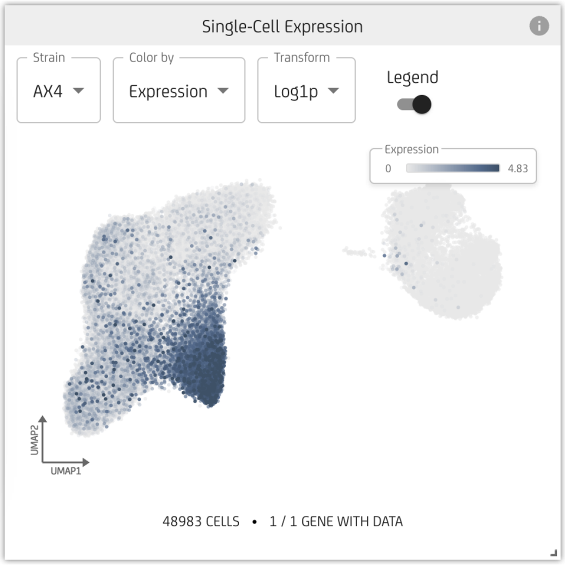


**B** AX4, coloured by *ecmA* expression

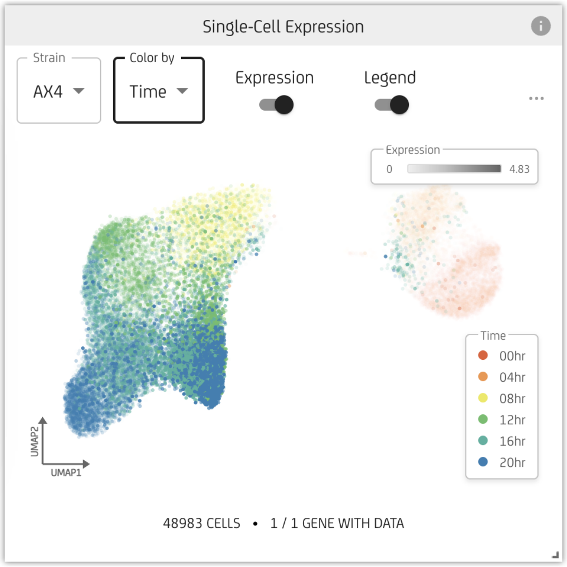


**C** AX4, coloured by developmental time

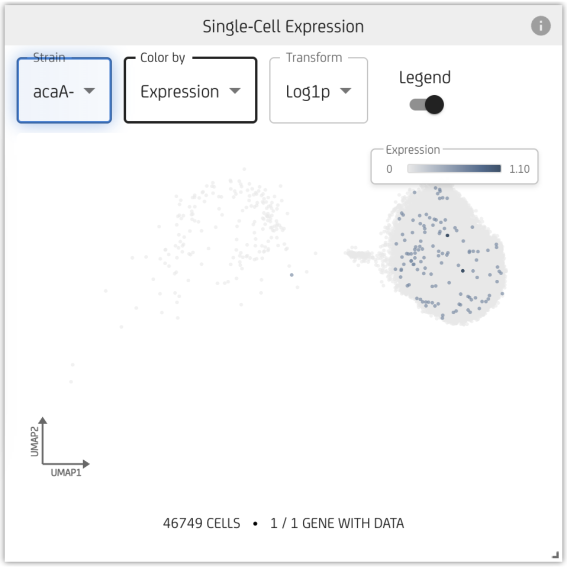


**D** *acaA*-, coloured by *ecmA* expression

Figure S1: **Opening workflow of the built-in tutorial.** (A) Select the developmental bulk dataset and inspect *ecmA*. (B) View the same gene in the AX4 single-cell map, where the tutorial identifies the *ecmA*-expressing population as committed prestalk cells. (C) Colour AX4 by collection time. (D) Switch to *acaA*- while retaining the reference embedding. Panels are direct outputs from tutorial steps 1–8.

### Quantitative check

We quantified the *ecmA* pattern used in the tutorial using the expression values served by dictyExpress. In AX4, mean *ecmA* abundance increased from 0.005 at 0 h to 11.023 at 20 h, while the fraction of cells with a value greater than zero increased from 0.4% to 59.6%. In *acaA*-, the mean at 20 h was 0.004 and 0.4% of cells had a value greater than zero. Thus, the numerical summary recapitulates the developmental increase and strain contrast seen in the interactive views.

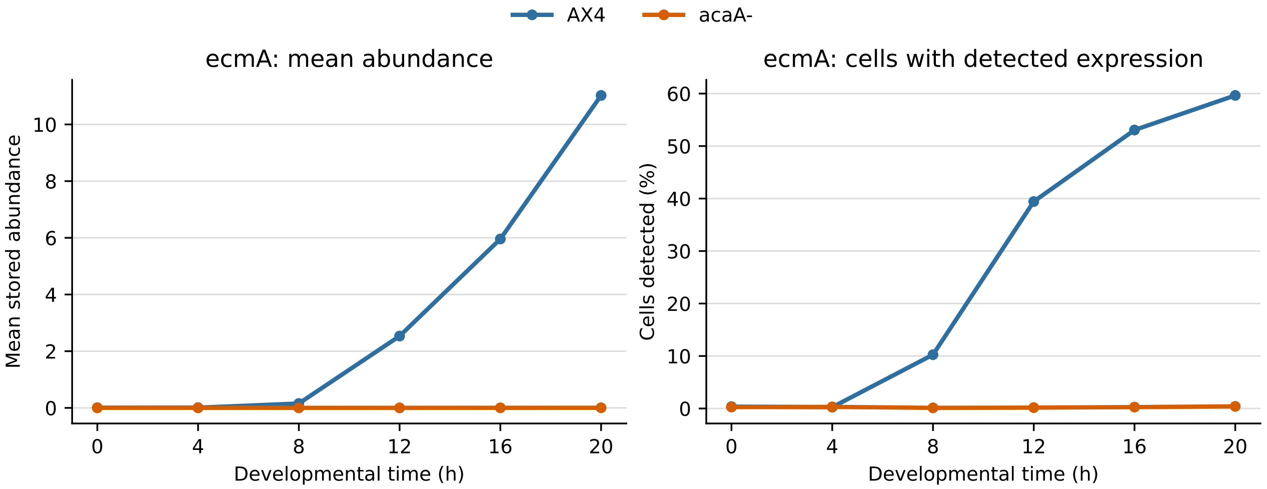


Figure S2: **Time-resolved *ecmA* summaries.** Mean stored abundance (left) and percentage of cells with a value greater than zero (right) are shown for AX4 and *acaA*-. These are descriptive cell-level summaries, not replicate-level estimates.

**Methods.** We used the processed single-cell matrices displayed by dictyExpress for AX4 (48,983 cells) and *acaA*- (46,749 cells), derived from GEO accession GSE305468. Cells were sampled at 0, 4, 8, 12, 16, and 20 h. For each strain and time point, we calculated the arithmetic mean of the stored *ecmA* abundance and the percentage of cells with a value greater than zero. No inferential tests were performed, and individual cells were not treated as independent biological replicates.

**Data source.** The processed data are available under GEO accession GSE305468. Katoh-Kurasawa M, Trnovec L, Lehmann P, et al. Early cAMP signaling orchestrates single-cell synchronicity throughout *Dictyostelium* development. *Commun Biol*. 2026;9:543. doi:10.1038/s42003-026-09806-5.
